# Assessment of carbon dioxide, carbon dioxide/oxygen, isoflurane and pentobarbital killing methods in rats

**DOI:** 10.1101/061002

**Authors:** Jessica M Chisholm, Daniel SJ Pang

**Affiliations:** Department of Veterinary Clinical and Diagnostic Sciences, University of Calgary, Alberta, Canada; Hotchkiss Brain Institute, University of Calgary, Alberta, Canada

## Abstract

**Background**:

Exposure to carbon dioxide (CO_2_) gas as a killing method is aversive and exposure to high concentrations likely to be painful. Bradycardia during exposure to CO_2_ is associated with nociception and pain. However, it is unclear if bradycardia occurs before loss of consciousness as this is variably defined in the literature. The objectives of this study were to explore the relationship between recumbency, loss of righting reflex (LORR) and a quiescent electromyograph as measures of loss of consciousness, and identify the onset of bradycardia in relation to these measures.

**Methods**:

Thirty-two adult, female Sprague-Dawley rats were instrumented with a telemetry device and randomly assigned to one of four killing methods (100% CO_2_, CO_2_ (70%)/O_2_ (30%), isoflurane (5%) and intraperitoneal pentobarbital (200 mg/kg). Time to achieve recumbency, LORR, quiescent electromyograph, isoelectric electrocorticograph, heart rate and apnea were recorded.

**Results**:

The general order of progression was recumbency, LORR, quiescent electromyograph, isoelectric electrocorticograph and apnea. Recumbency preceded LORR in the majority of animals (CO_2_; 7/8, CO_2_/O_2_; 8/8, isoflurane; 5/8, pentobarbital; 4/8). Bradycardia occurred before recumbency in the CO_2_ (p = 0.0002) and CO_2_/O_2_ (p = 0.005) groups, with a 50% reduction in heart rate compared to baseline. The slowest (time to apnea) and least consistent killing methods were CO_2_/O_2_ (1180 ± 658.1s) and pentobarbital (875 [239 to 4680]s).

**Conclusion**:

Bradycardia, and consequently nociception and pain, occurs before loss of consciousness during CO_2_ exposure. Pentobarbital displayed an unexpected lack of consistency, questioning its classification as an acceptable euthanasia method in rats.

## Introduction

The majority of laboratory rodents used in biomedical research are killed upon project completion. Ideally, the killing process is a “good death” (euthanasia), free from pain and distress.[1,2] The most recent Canadian Council on Animal Care (CCAC) and American Veterinary Medical Association (AVMA) euthanasia guidelines are broadly similar in their classification of killing methods.[1,2] Both guidelines consider CO_2_ to be “conditionally acceptable”/“acceptable with conditions” and overdose with intravenous or intra-peritoneal (IP) barbiturate as an acceptable method. In contrast, overdose with an inhalational anaesthetic agent (followed by a second method to ensure death after loss of consciousness) is considered acceptable by the CCAC and acceptable with conditions by the AVMA. Overdose with carbon dioxide (CO_2_) gas is a common killing method but exposure to low concentrations (< 20%) is aversive to rats and mice.[3–5] Despite this, CO_2_ remains popular as it is rapidly acting, simple to use, familiar, has a low risk of harm associated with human exposure and is effective for groups of animals. Exposure to the volatile anaesthetic agent, isoflurane, offers a refinement over CO_2_ by reducing, but not preventing, aversion in rats.[3,6] A less explored alternative, a mixture of CO_2_ and oxygen (CO_2_/ O_2_) has been associated with fewer signs of distress during exposure than CO_2_ alone, though results have been conflicting.[7–9]

When CO_2_ is employed, a gradual fill technique with displacement rates of between 10-30% of the chamber volume per minute (cv/min) are recommended to avoid pain resulting from exposure to high concentrations of CO_2_(>50%) prior to loss of consciousness.[1,2] The evidence for pain is from the human literature, with self-reports of nasal irritation and pain beginning at CO_2_ concentrations of > 35%. [10,11] Exposure to similar concentrations have been shown to activate nociceptors in rats[12–16] and result in reflex bradycardia.[17–19] Therefore, the observation of bradycardia during exposure to CO_2_ may serve as an indicator of nociception and potentially pain in rats.[20] If so, the timing of bradycardia in relation to loss of consciousness is critical to evaluating the presence of nociception or pain. However,there is confusion in the literature in how loss of consciousness is identified in rodents, leading to conflicting reports of the the occurrence of bradycardia before or after loss of consciousness.[20,21] There is currently no consensus over how to identify loss of consciousness in rats, with some studies relying on cessation of movement or recumbency.[20–23] This contrasts with experimental evidence suggesting that the appropriate surrogate measure of unconsciousness is loss of the righting reflex (LORR).[24]

Using 3 treatment groups, CO_2_, CO_2_/O_2_ and isoflurane, the aims of this study were:1. to compare three putative measures of loss of consciousness (recumbency, LORR and a quiescent electromyograph [EMG]) and examine the relationship of each to the presence of bradycardia and 2. to investigate the relationship between an isoelectric electrocorticograph (ECoG) and apnea as indicators of impending death. We hypothesised that bradycardia would precede the loss of righting reflex, indicating the possibility of pain prior to loss of consciousness and that the appearance of an isoelectric ECoG would be closely related to apnea. After initiating the project, a fourth treatment group, IP sodium pentobarbital (PB), was added as it was felt this would serve as a criterion standard for comparison.

## Materials and Methods

**Animals.** Experiments were performed at the University of Calgary following approval by the University of Calgary Health Science Animal Care Committee (protocol AC11-0044), which operates under the auspices of the CCAC.

Thirty-two female Sprague-Dawley rats (Health Science Centre Animal Resource Centre, Calgary, Alberta, Canada) weighing between 250 to 500 grams were used. Animals were housed in a 12h:12h light cycle (lights on at 0700h) and were group housed prior to instrumentation and singly housed afterwards, in micro-isolator rat cages (48 × 27 × 20cm [Ancare Corp., Worcester, MA, USA]). Fresh water and food (Prolab 2500 Rodent 5p14, Lab diet, PMI Nutrition International, St Louis MO, USA) were available ad libiFum. Plastic tubing (PVC pipe, provided by the Health Science Animal Resource Centre, Calgary, AB, Canada) wood shavings (Aspen chip, NEPCO, Warrensburg, NY, USA) and Nestlets (Nestlets nesting material, Ancare, Bellmore, New York, USA) were provided for bedding and enrichment. All experiments were performed between 1000h and 1600h.

### Treatment groups

Animals were block randomized (www.random.org) to one of four killing methods (n = 8 per group):CO_2_ (Praxair, Calgary, AB, Canada); exposure to 100% CO_2_ at a fill rate of 20% cv/min, isoflurane group; 5% isoflurane carried in oxygen at a fill rate of 20% cv/min until LORR, followed by stopping isoflurane administration and switching to 100% CO_2_ (30 % cv/min), CO_2_/O_2_; exposure to a mixture of 70% carbon dioxide and 30% oxygen at a fill rate of 20% cv/min and IP PB; (200 mg/kg, 240 mg/ml, Euthanyl,Bimedia MTC, Cambridge, ON, Canada).

### Telemetry instrumentation

Each rat was implanted with a radio transmitter (4ET-S2 Radio Transmitter Data Sciences International, St Paul, MN, USA) placed subcutaneously lateral to midline on the dorsum with leads for EMG, electrocardiography (ECG) and ECoG tunnelled subcutaneously to the central trapezius muscle of the neck (EMG), pectoral muscles (ECG) and skull (ECoG). Surgery for instrumentation was facilitated with general anesthesia as follows. General anesthesia was induced with isoflurane (5%) carried in oxygen (1 L/min), with rats placed singly in a perspex chamber. Following LORR the rat was moved to the surgical area and isoflurane (1.5-2%) delivered through a nose cone. Surgical sites were clipped and aseptically prepared and pre-emptive analgesia given. All animals received 0.1 ml (2 mg) of 2% lidocaine (diluted in 0.8 ml saline) as incisional line blocks, enrofloxican (50 mg/kg SC, 25 mg/ml, Baytril, Bayer, Toronto, ON, Canada), saline (4 ml, NaCl 0.9%, Baxter Corporation, Mississauga, Ontario, CA), buprenorphine 0.05 mg/ kg SC, every 8 hours (0.3 mg/ml Vetergesic, Champion Alstoe Animal Health, Whitby, ON, Canada) and meloxicam 1 mg/kg SC, every 24 hours (Metacam, Boehringer Ingelheim, Burlington, ON, Canada). Analgesics were continued for a minimum of 24 hours following surgery and pain assessed regularly (every 6-8 hours) by monitoring activity, posture, grooming and body weight. Antibiotics were continued for two days following the surgery. A minimum of 7 days passed before the experimental day.

For the experiment, animals were placed singly in a customised perspex chamber (25.5 (l) × 10 (w) × 12 (h) cm). The chamber had ports for gas entry and exit located on the short sides at opposite ends. The following physiological parameters were collected using commercial software (Data quest Advanced Research Technology version 4.3, Data Sciences International St. Paul, MN, USA):ECoG, EMG and ECG. The ECoG and EMG signals were sampled at 500 Hz with a 0 -100 Hz bandpass filter. The ECG signal was sampled at 1000 Hz with a 0 - 250 Hz bandpass filter. Baseline data were recorded over five minutes during exposure to room air. In the IP PB group, injections were given following baseline recording and the animal immediately returned to the recording chamber.

The following time points were recorded and compared to evaluate relationships between recumbency, LORR and muscle tone:baseline - recumbency, baseline - LORR, baseline - quiescent EMG. The times from baseline - isoelectric ECoG and baseline - apnea were used to investigate the relationship between an isoelectric ECoG and apnea. The overall speed of each method was assessed with the time between baseline - apnea.

Recumbency was defined as the moment when an animal’s body and head were in full contact with the chamber floor. The LORR was determined by manually rotating the chamber to place the animal on its back, assessing its ability to right itself. The onset of recumbency triggered the first assessment of LORR. LORR was confirmed if a rat could be turned on to its back for at least 10s. If LORR occurred at the first test, the same time was given for recumbency and LORR. An isoelecFc ECoG was identified by off-line visual inspection of the ECoG and defined as the waveform being within ± 0.025 mV of the x-axis, similar to the definition in humans (Fig. 1).[25]

**Figure 1:**
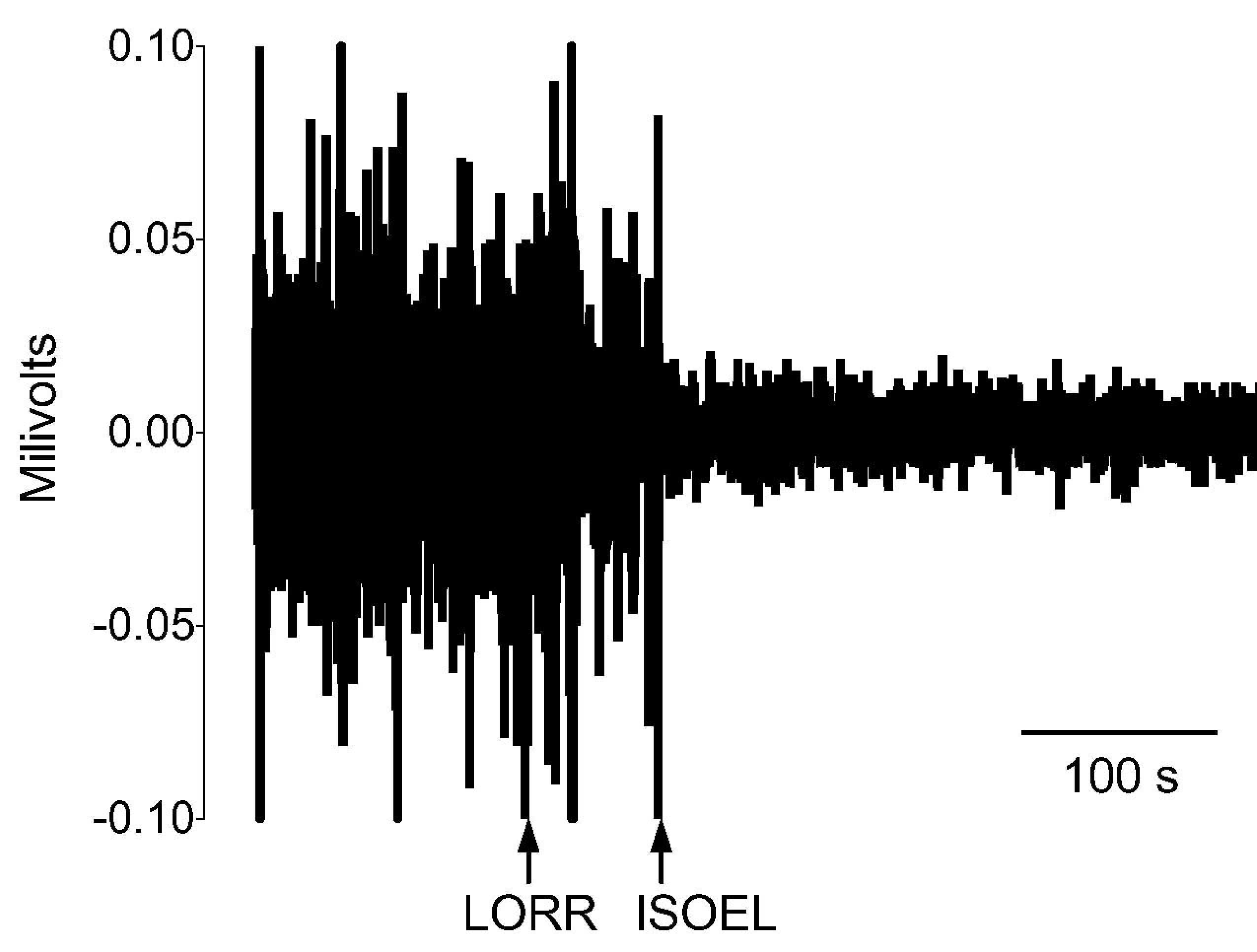
A representative example of the onset of an isoelectric electrocorticograph (ISOEL), occurring after loss of the righting reflex (LORR).

A quiescent EMG was determined by off-line visual inspection of the EMG and defined as the waveform being within XXX.

Heart rates were averaged over the 10 seconds immediately preceding each of the following times:end of baseline and occurrence of recumbency, LORR, isoelectric ECoG and apnea. Each rat was kept in the chamber until cardiac asystole was observed on the ECG. Death was confirmed by digital palpation of the thorax to confirm absence of a heart beat.

### Statistical analyses

Data were analysed with commercial software (Prism v7.0a, GraphPad Software Inc., La Jolla, CA, USA). Data were assessed for normality with a Shapiro-Wilk normality test. Differences between groups were compared with one-way ANOVA with a Tukey’s post hoc test. Heart rate data were analysed for differences within groups with a one-way ANOVA for repeated measures and a Dunnett’s post hoc test (comparison to baseline values). Where there was a significant change in heart rate between baseline and recumbency or LORR, unpaired t tests were used to compare heart rates between groups at these two time points. Pentobarbital data were handled separately and compared with the CO_2_ treatment group with either a Mann-Whitney test or unpaired t test, depending on distribution of the data. Coefficient of variation was calculated to provide an indication of data variability. A value of p < 0.05 was considered significant and 95% confidence intervals (95% CI) presented where available.

## Results

Data from the inhalational treatment groups were normally distributed. In the IP PB group heart rate data were normally distributed whereas time data were not.

### Recumbency precedes loss of righting reflex

Recumbency preceded LORR in 7/8 animals in the CO_2_ group (p = 0.30, 95%CI [-57.0, 14.5]), 8/8 animals in the CO_2_/O_2_ group (p = 0.16, 95%CI [-115.0, 16.7]) and 5/8 animals in the ISO group (p = 0.6, 95%CI [-82.0, 34.2]) with the time from recumbency to LORR ranging from 21.2-49.1 seconds (Table 1).

**Table 1:**
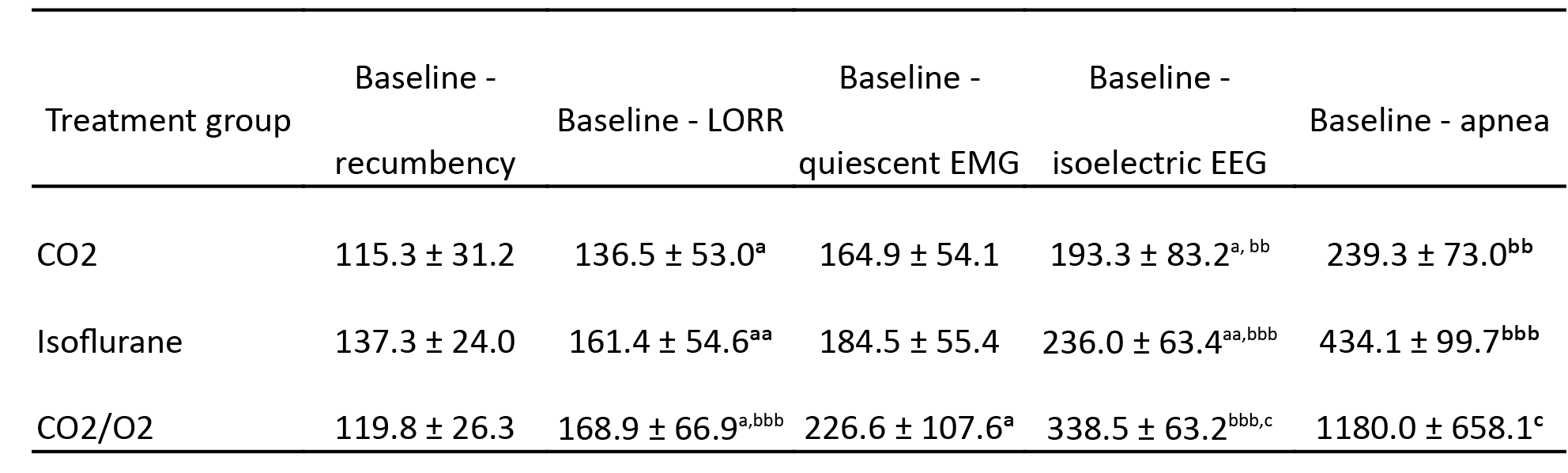
Same superscript letter denotes significant difference between time points within a group:single letter; p < 0.05, two letters; p < 0.01, three letters; p ≤ 0.001. Statistical comparisons were restricted to:recumbency vs. loss of righting reflex (LORR), LORR vs. quiescent electromyograph (EMG), LORR vs. isoelectric electrocorticograph (ECoG), isoelectric ECoG vs. apnea. See text and Figure 3 for results of between group comparisons. Data are mean ± SD.

| Treatment group | Baseline -<br>recumbency | Baseline - LORR | Baseline -<br>quiescent EMG | Baseline -<br>isoelectric EEG | Baseline - apnea |
| --- | --- | --- | --- | --- | --- |
| CO<sub>2</sub> | 115.3 ± 31.2 | 136.5 ± 53.0<sup>a</sup> | 164.9 ± 54.1 | 193.3 ± 83.2<sup>a, bb</sup> | 239.3 ± 73.0<sup>bb</sup> |
| Isoflurane | 137.3 ± 24.0 | 161.4 ± 54.6<sup>aa</sup> | 184.5 ± 55.4 | 236.0 ± 63.4<sup>aa,bbb</sup> | 434.1 ± 99.7<sup>bbb</sup> |
| CO<sub>2</sub>/O<sub>2</sub> | 119.8 ± 26.3 | 168.9 ± 66.9<sup>a,bbb</sup> | 226.6 ± 107.6<sup>a</sup> | 338.5 ± 63.2<sup>bbb,c</sup> | 1180.0 ± 658.1<sup>c</sup> |

There were no significant differences between inhalational treatment groups for the time from baseline to recumbency (CO_2_ vs. ISO, p = 0.26, 95%CI [-56.4, 12.4]; CO_2_vs CO_2_/O_2_, p = 0.94, 95% CI [-38.9, 29.9]; ISO vs CO_2_/O_2_, p = 0.42, 95% CI [-16.9, 51.9], Table 1). Similarly, there were no significant differences between inhalational treatment groups from baseline to LORR (CO_2_ vs. ISO, p = 0.68, 95%CI [-98.6, 48.8]; CO_2_ vs CO_2_/O_2_, p = 0.52, 95% CI [-106.1, 41.3]; ISO vs CO_2_/O_2_, p = 0.96, 95% CI [-81.2, 66.2]). LORR preceded EMG quiescence in all animals in the CO_2_/O_2_ treatment group, with one animal in the CO_2_ group and two animals in the ISO group exhibiting EMG quiescence prior to LORR. The mean delay between LORR and EMG quiescence ranged from 23.1 seconds for ISO to 57.8 seconds for CO_2_/O_2_, with a significant delay in the CO_2_/O_2_ group (Table 1). There were no significant differences between inhalational treatment groups between LORR and a quiescent EMG (CO_2_ vs. ISO, p = 0.95, 95%CI [-37.9, 48.4]; CO_2_ vs CO_2_/O_2_, p = 0.22, 95% CI [-72.5, 13.8]; ISO vs CO_2_/O_2_, p = 0.13, 95% CI [-77.8, 8.5]).

PB did not differ significantly from the CO_2_ group in the phases between baseline and recumbency (p = 0.43) or baseline and LORR (p = 0.12, Table 2), with recumbency preceding LORR in 4/8 animals.

However, in contrast to the inhalational treatment groups, EMG quiescence preceded LORR in 7/8 animals. This early onset of EMG quiescence was significantly faster than the CO_2_ group (p = 0.004)

**Table 2:**
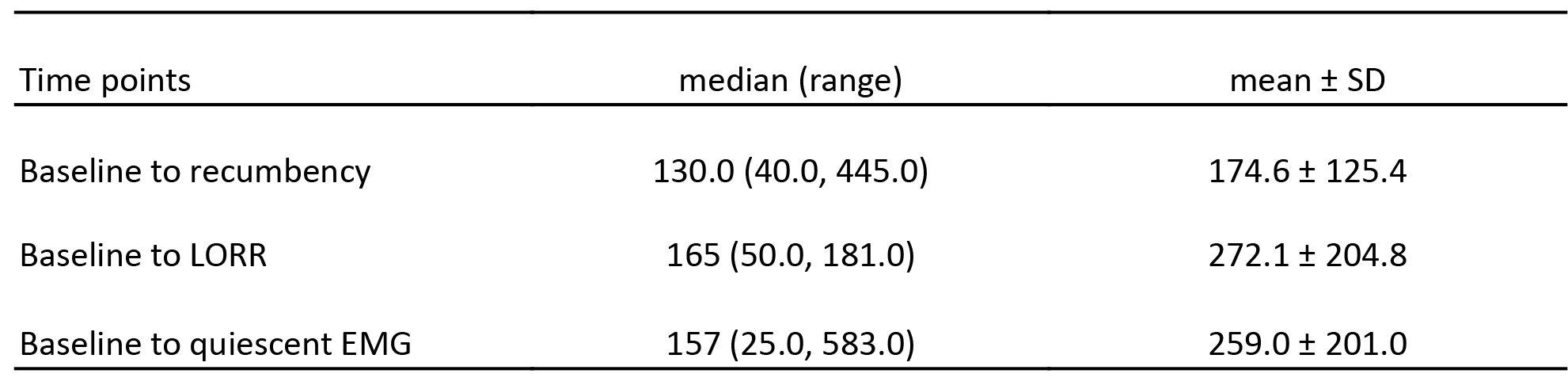
Recorded times for recumbency, loss of righting reflex (LORR) and electromyography (EMG) quiescence in the pentobarbital treatment group. Statistical comparisons with the CO_2_ treatment group were performed with median (range) data; mean ± SD are provided for completeness.

| Time points | median (range) | mean ± SD |
| --- | --- | --- |
| Baseline to recumbency | 130.0 (40.0, 445.0) | 174.6 ± 125.4 |
| Baseline to LORR | 165 (50.0, 181.0) | 272.1 ± 204.8 |
| Baseline to quiescent EMG | 157 (25.0, 583.0) | 259.0 ± 201.0 |

Bradycardia precedes both recumbency and loss of righting reflex

Heart rates did not differ between treatment groups at baseline (CO_2_ vs. CO_2_/O_2_, p = 0.58, 95%CI [-27.9, 64.9]; CO_2_ vs. isoflurane, p = 0.44, 95% CI [-69.5, 23.3]; CO_2_/O_2_ vs. isoflurane, p = 0.08, 95%CI [-4.8, 88.0]; CO_2_ vs. PB, p = 0.52, 95% CI [-26.2, 49.4]), with average values ranging from 396 to 438 beats per minute (Fig. 2, Table 3).

**Table 3:**
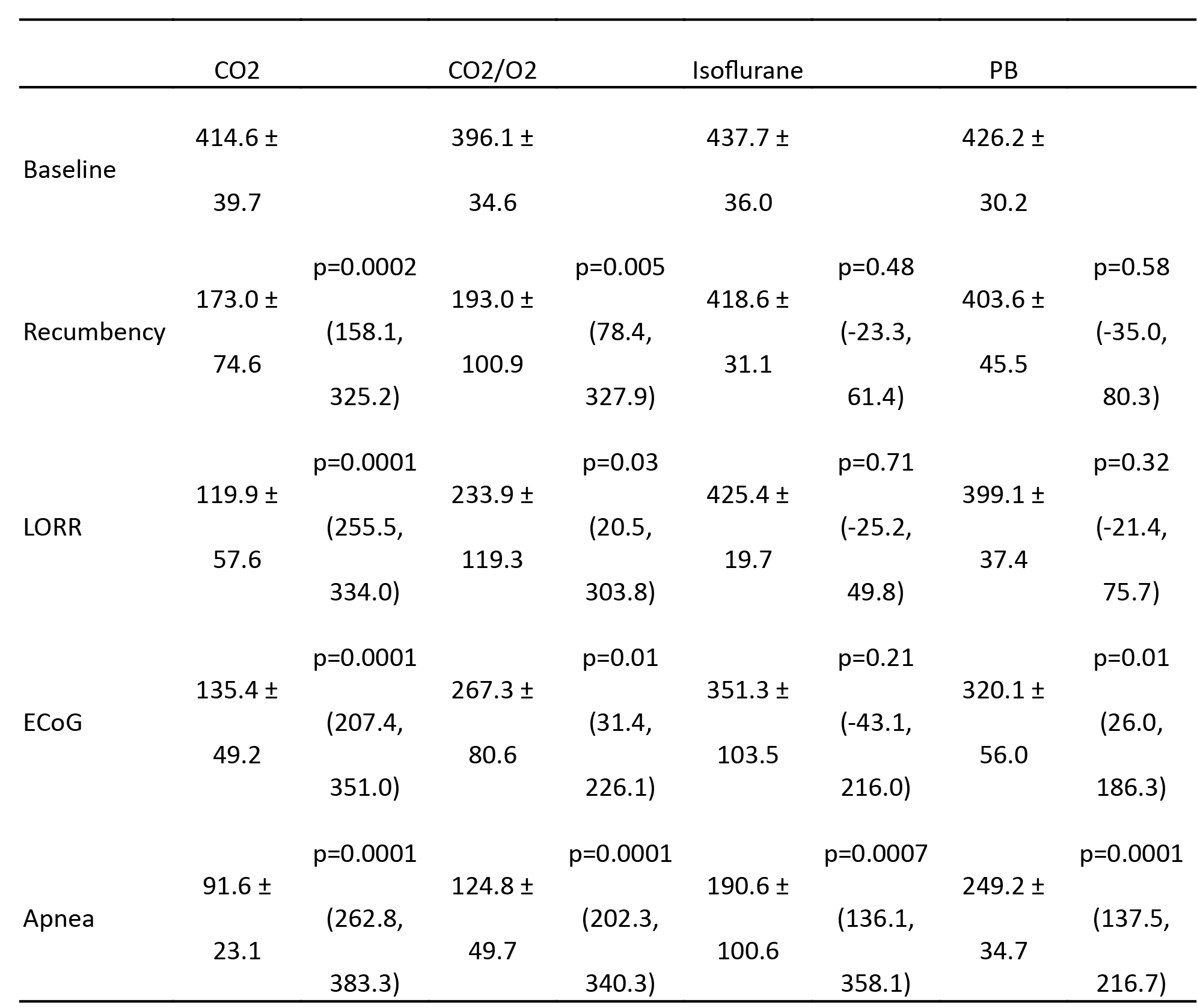
Heart rates (beats per minute) recorded at different time points in treatment groups. PB = pentobarbital. LORR = loss of righting reflex. ECoG = isoelectric electrocorticograph. p values represent within group comparisons to baseline. See text and Figure 2 for results of between group comparisons. Data are mean ± SD.

Bradycardia prior to loss of the righting reflex only occurred in the CO_2_and CO_2_/O_2_ groups (Fig. 2, Table 3) with an average decrease of 58.3% and 51.3%, respectively. In the isoflurane and PB treatment groups, bradycardia appeared at or after the onset of an isoelectric ECoG (Table 3). At recumbency, the bradycardia observed in the CO_2_ and CO_2_/O_2_ groups was significantly lower than the isoflurane group (isoflurane vs. CO_2_, 95% CI [-339.7, -151.5]; isoflurane vs CO_2_/O_2_, 95% CI [-319.7, -131.5]; p < 0.0001 both comparisons, Fig. 2). There was no significant difference between CO_2_ and CO_2_/O_2_ groups at recumbency (p = 0.85, 95% CI [-114.1, 74.1]) but heart rate was significantly higher (approximately double) in the CO_2_/O_2_ group at the LORR (p = 0.008, 95% CI [-202.3, -25.7], Fig 2). Both CO_2_ and CO_2_/O_2_ groups had significantly lower rates than the isoflurane group (isoflurane vs. CO_2_, 95% CI [-393.8, -217.2]; isoflurane vs CO_2_/O_2_, 95% CI [-279.8, -103.2]; p < 0.0001 both comparisons). Heart rates in all groups converged at the point of apnea (Table 3).

**Figure 2:**
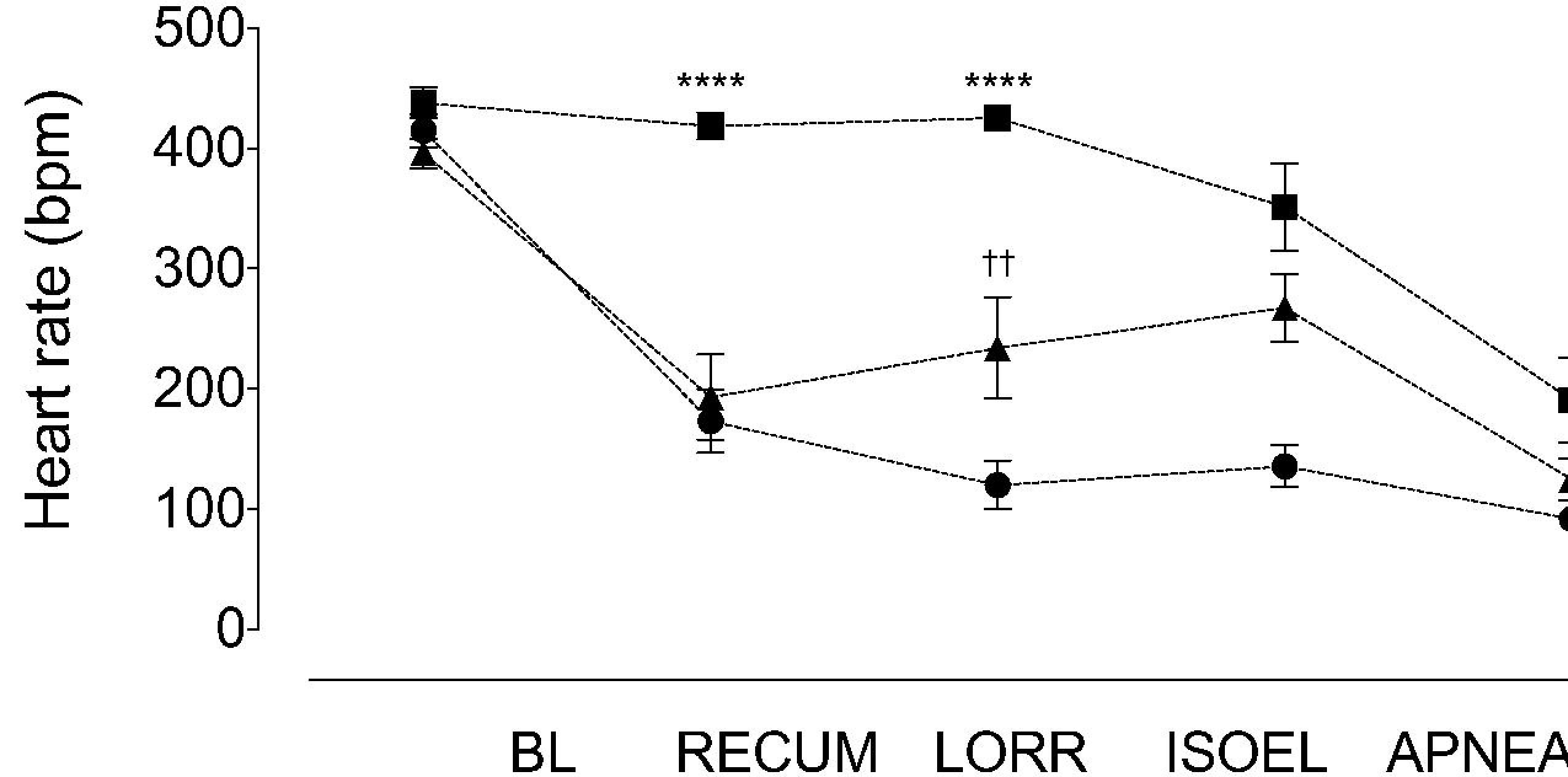
Heart rates in the carbon dioxide (circles) and carbon dioxide-oxygen (triangles) treatment groups decrease significantly compared to the isoflurane group (squares) at recumbency (RECUMB, **** p < 0.0001, both comparisons) and loss of the righting reflex (LORR, **** p < 0.0001, both comparisons). At LORR, heart rates are significantly increased in the carbon dioxide-oxygen group compared with the carbon dioxide group (†† p = 0.008). ISOEL, isoelectric electrocorticograph. Data are mean ± SEM. Isoelectric ECoG occurs after loss of righting reflex and precedes apnea

An isoelectric ECoG occurred after LORR in all animals, representing an increasing depth of anaesthesia (Fig. 3A). The onset of an isoelectric ECoG was shortest in the CO_2_ group (Table 1). This was not significantly different from the isoflurane group (p = 0.73, 95% CI [-76.6, 40.9]) and occurred sooner than in the CO_2_/O_2_ group (169.6 ± 50.2 seconds, p = 0.0002, 95% CI [-171.6, -54.1]). Onset of an isoelectric ECoG was also earlier in the isoflurane group compared with the CO_2_/O_2_ group (p = 0.002, 95% CI [-153.8, -36.3]). The PB group did not differ from the CO_2_ group, but exhibited considerable data variability (p = 0.06, 101 [25.0 to 2342.0] seconds).

Apnea occurred after an isoelectric ECoG in all cases (Fig. 3B). This period was shortest for the CO_2_ group (Table 1) and was significantly faster compared with the CO_2_/O_2_ group (p = 0.002, 95% CI [-1288, 302.6]),but not the isoflurane group (p = 0.72, 95% CI [-644.7, 340.4]). This time course was also shorter in the isoflurane compared with the CO_2_/O_2_ group (p = 0.009, 95% CI [-1136.0, 150.5]). The PB group did not differ from the CO_2_ group, but again displayed large data variability (287.5 [4.0 to 4200.0 seconds], p = 0.07).

The Fme course for the entire observation period (from baseline until apnea) was fastest in the CO_2_ and ISO groups (Fig. 3C, Table 1). Though there was no significant difference between the CO_2_ and ISO group (p = 0.61, 95% CI [-669.0, 304.0]), the average time to apnea in the CO_2_ group (239.3 ± 73.0 seconds) was approximately half that of the ISO group (434.1 ± 99.7 seconds). The source of the increased time to apnea in the ISO group resulted from a four fold increase in average time between isoelectric ECoG and apnea compared to the CO_2_ group (Fig. 3B, Table 1). Both CO_2_ and isoflurane treatment groups reached apnea faster than the CO_2_/O_2_ group (vs. CO_2_, p = 0.0003, 95% CI [-1415.0, -441.0]; vs. ISO, p = 0.003, 95% CI [-1232.0, -259.0]). Time to apnea was faster in the CO_2_ group than the PB group (p = 0.005, 875 [239 to 4680] seconds). The most consistent killing methods, with the lowest coefficients of variation, were CO_2_ (26.9%) and ISO (23.0%), followed by CO_2_/O_2_ (55.8%) and PB (114.1%). In the PB treatment group, three rats contributed to substantial variability in the data set, as a result of suspected misinjection.

**Figure 3:**
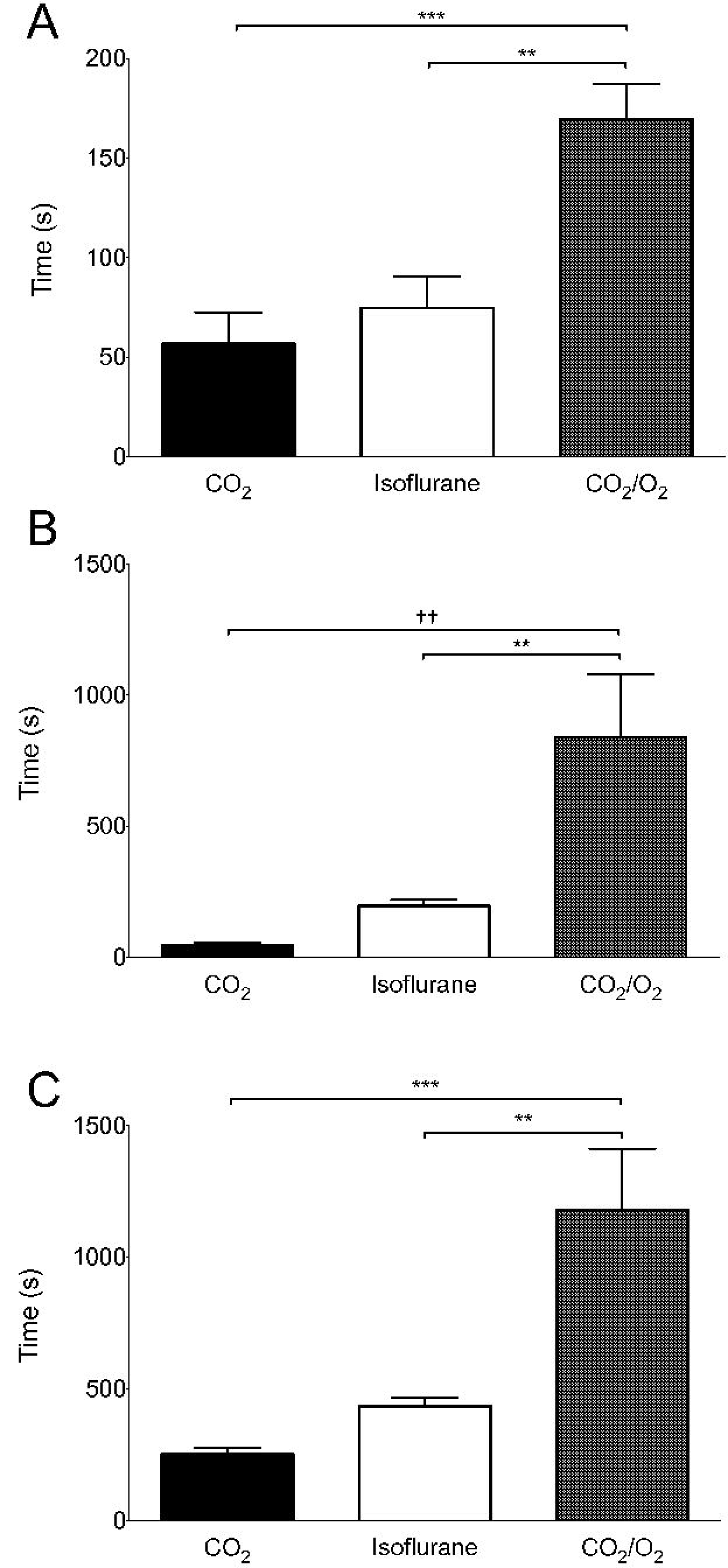
Time periods during which differences between treatment groups emerged. A:Time from loss of the righting reflex until an isoelectric electrocorticograph. *** p = 0.0002, ** p = 0.002. B:Time from an isoelectric electrocorticograph until apnea. ++ p = 0.002, ** p = 0.01. C:Time from baseline until apnea. ** p = 0.003, *** p = 0.0003. CO_2_, carbon dioxide. CO_2_/O_2_, carbon dioxide/oxygen. Data are mean ± SEM.

## Discussion

In evaluating euthanasia methods the AVMA Guidelines for the Euthanasia of Animals include assessment of the following criteria:the “time required to induce loss of consciousness”, “reliability” and the “ability to induce loss of consciousness and death with a minimum of pain and distress”.[1] Our data provide insight on the time to loss of consciousness and reliability of the studied methods, allowing comment on the potential for pain and distress.

We have shown that:1. LORR and recumbency occur at different times, indicating that recumbency is not an accurate indicator of loss of consciousness, 2. bradycardia occurs in response to exposure to carbon dioxide gas both with and without supplemental oxygen and that bradycardia precedes LORR, 3. euthanasia with a gradual fill carbon dioxide technique is the fastest of the methods studied to achieve apnea but the time to LORR did not differ between carbon dioxide and isoflurane. The addition of supplemental oxygen during carbon dioxide euthanasia substantially increases Fme to apnea and 4. considerable variability is associated with both CO_2_/O_2_ and IP PB methods, questioning the classification of IP PB as an acceptable euthanasia method.[1,2]

There is a strong positive correlation between LORR in rodents and unconsciousness in humans, suggesting that LORR is an appropriate proxy for loss of consciousness in rats.[24] The onset of LORR equates to a light plane of anaesthesia, insufficient to prevent movement in response to a noxious stimulus, approximating MAC_awake_ in humans, where MAC is the minimum alveolar concentration of an inhalational anaesthetic agent which prevents gross, purposeful movement in response to a supramaximal noxious stimulus in an individual (or 50% of a study population).[26] And MAC_awake_ is the lower concentration of anaesthetic, approximately 50% of MAC, when an individual (or 50% of a study population) can provide a verbal response to a command.[27]

Recumbency preceded LORR in the majority of animals studied. This suggests that previous investigations which used recumbency as a proxy for loss of consciousness underestimated the speed to reach loss of consciousness.[20–23] As the Fme between initiation of the killing process and unconsciousness is a critical period when pain may be perceived, the reliance on recumbency has implications for the assessment of welfare of killing methods. In this study, the mean Fme to achieve recumbency in the CO_2_ group of 115 seconds, is similar to that previously reported where gradual fill techniques were used.[14,20,21,23]

Moody et al (2015) suggested a more conservative indicator of unconsciousness, an absent pedal withdrawal reflex.[28] This undoubtedly reduces the risk that an animal may be conscious during exposure to a noxious stimulus, a valid consideration when deciding to expose an animal to such a stimulus (e.g. high concentration CO_2_, surgery). However, the literature suggests that movement can occur when an animal (or person) is unconscious as the concentration of anaesthetic required to induce loss of consciousness is lower than that required to abolish movement.[27,29–31]

Residual muscle activity beyond loss of consciousness was reflected in the time to achieve a quiescent EMG exceeding that required for LORR. Hewett et al (1993) observed increased muscle tonicity during exposure to high concentrations (>90%, pre-fill) of CO_2_ and spontaneous muscle activity can continue after death.[21,32] Together, this indicates that appearance of a quiescent EMG is an insensitive indicator of unconsciousness.

An isoelectric ECoG represents depressed cortical function, beyond that typically observed with therapeutic doses of anaesthetic and analgesic drugs.[33] However, the presence of an isoelectric ECoG alone is insufficient to confirm death.[34–36] Our results show that the time between onset of the isoelectric EEG and apnea varied considerably between treatment groups, taking up to 14 minutes in the CO_2_/O_2_ group in contrast to approximately 45 seconds in the CO_2_ group. The prolonged Fme to achieve an isoelectric ECoG in the isoflurane and CO_2_/O_2_ treatment groups suggests that providing O_2_ may delay its onset and the Fme to apnea.

The potential benefit of using a mixture of CO_2_ and O_2_ for euthanasia is controversial.[7–9] Coenen et al. (1995) reported that the combination of oxygen and carbon dioxide, delivered at a high chamber fill rate (188% cv/min, 2:1 CO_2_:O_2_ ratio) prevented gasping when compared with carbon dioxide alone.[7] In contrast, Iwarsson and Rehbinder (1993) observed laboured breathing and “uneasiness” during exposure to a chamber pre-filled with carbon dioxide (80%) and oxygen (20%).[8] The combination of CO_2_and O_2_ has a modest effect on reducing aversion to the gas mixture in comparison to CO_2_ alone.[9] These studies also reported a prolonged time to death with CO_2_/O_2_ compared with CO_2_ alone despite the rapid rate of exposure. This slowing of the killing process reflects our observations that, when compared with CO_2_ alone, the time from LORR to apnea was 10 times longer in the CO_2_/O_2_ group. Up to the point of LORR there was no significant difference between these two groups.

Given the conflicting reports of behaviours associated with respiratory distress, a prudent response to available evidence which takes in to account the AVMA guidelines for evaluating killing methods is to avoid the addition of O_2_ to CO_2_.[1]

In humans, nasal exposure to CO_2_ concentrations of approximately 35% are reported as moderately irritating, with irritation increasing as CO_2_ concentrations increase.[10,11] At similar concentrations, conjunctival and corneal exposure to CO_2_ result in stinging and burning sensations.[37,38] The onset of pain (nasal and ocular) begins at concentrations of CO_2_ of approximately 40%.[13,14] and this corresponds to nociceptor activation in rats beginning at a CO_2_ concentration of around 40%.[12,15,16] The perception of pain occurs at CO_2_ concentrations slightly (< 10%) above that of nociceptor activation in humans.[39]

Exposure of the nasal mucosa to CO_2_ in rats at concentrations associated with irritation and pain in humans results in a reflex bradycardia, mediated through the vagal nerve via baro-and chemoreflexes. [17,19] Our finding that bradycardia occurs prior to LORR contrasts with those of Hawkins et al. (2006), when bradycardia was observed approximately 120 seconds after recumbency.[20] Similar to our findings, two studies that recorded recumbency, but not LORR, observed bradycardia near the onset of recumbency.[7,21] Furthermore, the gas flow rates used (14 and 22% cv/min) and measurement of chamber CO_2_ indicated that bradycardia occurred at a concentration of CO_2_ lower than the 100% reported by Yavari et al. (1996).[19] Unfortunately, we did not record CO_2_ concentration in our testing chamber.

The variability observed in the PB group was considerably worse than expected and suspected to result from misinjection. Unfortunately, necropsy examinations were not performed and the PB solution used did not include a coloured dye. Intraperitoneal misinjection has been previously documented in rats, reporting rates of 6-20% by trained, experienced personnel.[40–42] There are several potential sites for inadvertent placement of the injectate, including intra-abdominal fat, the abdominal wall, subcutaneous space, retroperitoneal space and viscera.[40–42] Of these, placement in to viscera, predominantly the cecum, appears the most common site of misinjection.[42] The cecum in rats is usually located in the caudal left quadrant of the abdominal cavity. However, its location varies considerably, lying in the middle of the caudal region of the abdomen in 10-18% of rats and in the caudal right quadrant in 16-30%.[41]

Strategies to reduce misinjection rates include using a two person injection technique (as in this study), minimising the distance the needle is inserted in to the abdominal cavity and performing the injection with the head lowered below the level of the caudal abdomen.[41] However, the efficacy of these strategies is largely unproven.

Though the incidence of misinjection could not be determined in our study, the high coefficient of variation and wide variability observed for the total observation period (baseline to apnea) raises the index of suspicion that misinjection occurred. Concerningly, the time to recumbency and LORR did not differ significantly compared to the CO_2_ group, with the delay to apnea occurring after these end points. This highlights the importance of confirming death.[1,2]

The possibility of nociception or pain associated with administering IP PB has been identified by two studies, using behavioural and molecular evidence.[43,44] Where misinjection delays the Fme to death, it is unknown if pain may be present in animals unable to show behavioural changes. The observed variability when using IP PB suggests that its current classification as an “acceptable” needs re-evaluation to account for route of administration.[1,2]

This study had several limitations. We were unable to determine an accurate time of death as animals were left undisturbed in the test chamber until all cardiac electrical activity had ceased. It is highly likely that pulseless electrical activity would have been present, which without concurrent arterial blood pressure recording, prevents accurate determination of death. Consequently, apnea was used as the study end-point. The time between apnea and loss of pulsatile blood flow was previously reported as approximately one minute using a 22% cv/min gradual fill technique with 100% CO_2_.[21] The time from baseline to apnea in the isoflurane group could have been shortened by increasing the flow rate of CO_2_ gas after LORR occurred. In doing so, it is likely that the time to produce apnea would have been closer to that of the CO_2_ group. This study was not designed to explore the cause(s) of the inconsistent results seen in the PB group. Further work is necessary to determine if intra-peritoneal overdose with PB can be improved. Our results are limited to the strain and sex studied.

## Conclusions

The onset of recumbency is an inaccurate indicator of loss of consciousness in rats exposed to CO_2_ CO_2_/O_2_ and isoflurane, underestimating the Fme when pain may be perceived and during which there is also limited motor function. Bradycardia occurred in both COs_2_-containing groups prior to LORR. As bradycardia in rats exposed to CO_2_ occurs at a concentration reported as painful in humans, this highlights the possibility of rats experiencing pain prior to loss of consciousness.

Overdose with intraperitoneal PB did not produce consistent results, leading to the possibility of prolonged euthanasia times. This lack of reliability questions its classification as an acceptable euthanasia method.

## Acknowledgments

The authors wish to thank Dr Aleks Krajacic and Dr Deb De Rantere for technical assistance and Dr Darrel Florence for insightful discussions during project planning.400

## References

1. Association AVM (2013) AVMA guidelines for the euthanasia of animals:2013 edition. Available:www.avma.org/KB/Policies/Documents/euthanasia.pdf

2. Charbonneau R, Niel L, Olfert E, von Keyserlingk M, Griffin G (2010) CCAC guidelines on:euthanasia of animals used in science. Available:www.ccac.ca/en_/standards/guidelines

3. Leach MC, Bowell VA, Allan TF, Morton DB (2002) Aversion to gaseous euthanasia agents in rats and mice. Comp Med 52:249–257.

4. Niel L, Weary DM (2007) Rats avoid exposure to carbon dioxide and argon. Appl Anim Behav Sci 107:100–109.

5. Niel L, Stewart SA, Weary DM (2008) Effect of flow rate on aversion to gradual-fill carbon dioxide exposure in rats. Appl Anim Behav Sci 109:77–84.

6. Makowska IJ, Weary DM (2009) Rat aversion to induction with inhalant anaesthetics. Appl Anim Behav Sci 119:229–235.

7. Coenen AM, Drinkenburg WH, Hoenderken R, van Luijtelaar EL (1995) Carbon dioxide euthanasia in rats:oxygen supplementation minimizes signs of agitation and asphyxia. Lab Anim 29:262–268.

8. Iwarsson K, Rehbinder C (1993 A study of different euthanasia techniques in guinea pigs, rats, and mice. Animal response and postmortem findings. Scand J Lab Anim Sci 20:191–205.

9. Kirkden RD, Niel L, Stewart SA, Weary DM (2008) Gas killing of rats:the effect of supplemental oxygen on aversion to carbon dioxide. Anim Welfare 17:79–87.

10. Shusterman D, Avila PC (2003) Real-time monitoring of nasal mucosal pH during carbon dioxide stimulation:implications for stimulus dynamics. Chem Senses 28:595–601.

11. Wise PM, Wysocki CJ, Radil T (2003) Time-intensity ratings of nasal irritation from carbon dioxide. Chem Senses 28:751–760.

12. Anton F, Peppel P, Euchner I, Handwerker HO (1991) Controlled noxious chemical stimulation:responses of rat trigeminal brainstem neurones to CO_2_ pulses applied to the nasal mucosa. Neurosci Lett 123:208–211.

13. Anton F, Euchner I, Handwerker HO (1992) Psychophysical examination of pain induced by defined CO_2_ pulses applied to the nasal mucosa. Pain 49:53–60.

14. Danneman PJ, Stein S, Walshaw SO (1997) Humane and practical implications of using carbon dioxide mixed with oxygen for anesthesia or euthanasia of rats. Lab Anim Sci 47:376–385.

15. Peppel P, Anton F (1993) Responses of rat medullary dorsal horn neurons following intranasal noxious chemical stimulation:effects of stimulus intensity, duration, and interstimulus interval. J Neurophysiol 70:2260–2275.

16. Thurauf N, Friedel I, Hummel C, Kobal G (1991) The mucosal potential elicited by noxious chemical stimuli with CO_2_ in rats:is it a peripheral nociceptive event? Neurosci Lett 128:297–300.

17. Kobayashi M, Cheng ZB, Nosaka S (1999) Inhibition of baroreflex vagal bradycardia by nasal stimulation in rats. Am J Physiol 276:H176–84.

18. Kobayashi M, Majima Y (2004) Target site of inhibition of baroreflex vagal bradycardia by nasal stimulation. Brain Res 1009:137–146.

19. Yavari P, McCulloch PF, Panneton WM (1996) Trigeminally-mediated alteration of cardiorespiratory rhythms during nasal application of carbon dioxide in the rat. J Auton Nerv Syst 61:195–200.

20. Hawkins P, Playle L, Golledge H, Leach MC, Banzett R et al. (2006) Newcastle consensus meeting on carbon dioxide euthanasiaoflaboratoryanimals. Available:nc3rs.org.uk/euthanasia via the Internet.

21. Smith W, Harrap SB (1997) Behavioural and cardiovascular responses of rats to euthanasia using carbon dioxide gas. Lab Anim 31:337–346.

22. Correia R, Pereira A, Gabriel J, Antunes L (2015) Anaesthesia induction in small mammal’s using an instrumented anaesthetic chamber. IEEE 4th Portugese BioEngineering Meeting

23. Niel L, Weary DM (2006) Beahvioural responses of rats to gradual-fill carbon dioxide euthanasia and reduced oxygen concentrations. Appl Anim Behav Sci 100:295–308.

24. Franks NP (2008) General anaesthesia:from molecular targets to neuronal pathways of sleep and arousal. Nat Rev Neuroscis 9:370–386.

25. Azabou E, Fischer C, Mauguiere F, Vaugier I, Annane D et al. (2016) Prospective Cohort Study Evaluating the Prognostic Value of Simple EEG Parameters in Postanoxic Coma. Clin EEG Neurosci 47:75–82.

26. Eger EIn, Saidman LJ, Brandstater B (1965) Minimum alveolar anesthetic concentration:a standard of anesthetic potency. Anesthesiology 26:756–763.

27. Stoelting RK, Longnecker DE, Eger EIn (1970) Minimum alveolar concentrations in man on awakening from methoxyflurane, halothane, ether and fluroxene anesthesia:MAC awake. Anesthesiology 33:5–9.

28. Moody CM, Makowska IJ, Weary DM (2015) Testing three measures of mouse insensibility following induction with isoflurane or carbon dioxide gas for a more humane euthanasia. Appl Anim Behav Sci 163:183–187.

29. Antognini JF, Schwartz K (1993) Exaggerated anesthetic requirements in the preferentially anesthetized brain. Anesthesiology 79:1244–1249.

30. Antognini JF, Carstens E, Atherley R (2002) Does the immobilizing effect of thiopental in brain exceed that of halothane? Anesthesiology 96:980–986.

31. Antognini JF, Barter L, Carstens E (2005) Overview movement as an index of anesthetic depth in humans and experimental animals. Comp Med 55:413–418.

32. Hewett TA, Kovacs MS, Artwohl JE, Bennett BT (1993) A comparison of euthanasia methods in rats, using carbon dioxide in prefilled and fixed flow rate filled chambers. Lab Anim Sci 43:579–582.

33. Powner DJ ( 1976) Drug-associated isoelectric EEGs. A hazard in brain-death certification. JAMA 236:1123.

34. Citerio G, Crippa IA, Bronco A, Vargiolu A, Smith M (2014) Variability in brain death determination in europe:looking for a solution. Neurocrit Care 21:376–382.

35. Kimura J, Gerber HW, McCormick WF (1968) The isoelectric electroencephalogram. Significance in establishing death in patients maintained on mechanical respirators. Arch Intern Med 121:511–517.

36. Shemie SD, Hornby L, Baker A, Teitelbaum J, Torrance S et al. (2014) International guideline development for the determination of death. Intensive Care Med 40:788–797.

37. Chen X, Gallar J, Pozo MA, Baeza M, Belmonte C (1995) CO_2_ stimulation of the cornea:a comparison between human sensation and nerve activity in polymodal nociceptive afferents of the cat. Eur J Neurosci 7:1154–1163.

38. Feng Y, Simpson TL (2003) Nociceptive sensation and sensitivity evoked from human cornea and conjunctiva stimulated by CO_2_. Invest Ophthalmol Vis Sci 44:529–532.

39. Thurauf N, Gunther M, Pauli E, Kobal G (2002) Sensitivity of the negative mucosal potential to the trigeminal target stimulus CO(2). Brain Res 942:79–86.

40. Ballard T (2009) Intraperitoneal route of administration - how accurate is thistechnique. Animal Technology and Welfare 8:17–18.

41. Coria-Avila GA, Gavrila AM, Menard S, Ismail N, Pfaus JG (2007) Cecum location in rats and the implications for intraperitoneal injections. Lab Anim (NY) 36:25–30.

42. Lewis RE, Kunz AL, Bell RE (1966) Error of intraperitoneal injections in rats. Lab Anim Care 16:505–509.

43. Ambrose N, Wadham J, Morton D (1999) Refinement of euthanasia. 3rd World Congress on Alternatives and Animal Use in the Life Sciences 1159–1170.

44. Svendsen O, Kok L, Lauritzen B (2007) Nociception after intraperitoneal injection of a sodium pentobarbitone formulation with and without lidocaine in rats quantified by expression of neuronal c-fos in the spinal cord-a preliminary study. Lab Anim 41:197–203.

